# Endocrine autoimmune disease as a fragility of immune-surveillance against hypersecreting mutants

**DOI:** 10.1101/845750

**Authors:** Yael Korem Kohanim, Avichai Tendler, Avi Mayo, Nir Friedman, Uri Alon

## Abstract

Many endocrine organs show prevalent autoimmune diseases (AID) such as type-1-diabetes and Hashimoto’s-thyroiditis. The fundamental origins of these diseases is unclear. Here we address AID from the viewpoint of feedback control. Endocrine tissues maintain their mass by feedback-loops that balance cell proliferation and removal according to input signals related to the hormone function. Such feedback is unstable to mutant cells that mis-sense the signal, and therefore hyper-proliferate and hyper-secrete the hormone. We hypothesize that in order to prevent these mutants from expanding, each organ has a dedicated ‘autoimmune surveillance of hyper-secreting mutants’ (ASHM), in which hyper-secreting cells are preferentially eliminated, at the cost of a fragility to AID. ASHM correctly predicts the identity of the self-antigens and the presence of T-cells against these self-antigens in healthy individuals. It offers a predictive theory for which tissues get frequent AID, and which do not and instead show frequent mutant-expansion disease (e.g. hyperparathyroidism).

## Introduction

Autoimmune diseases can be classified into systemic disease, such as lupus which attacks many different organs, and organ specific diseases, such as type-1 diabetes. The organ specific diseases are usually based on T-cell attack of specific tissues.

Many organ-specific diseases occur in endocrine and exocrine organs. In type-1 diabetes, for example, T-cells target and kill pancreatic beta cells. Other common T-cell-based autoimmune diseases include Hashimoto’s disease of the thyroid, Addison’s disease of the adrenal cortex, and vitiligo in which skin melanocytes are attacked. These autoimmune diseases have a young age of onset, and their prevalence can exceed 0.1%-1% of the population. Interestingly, some endocrine organs very rarely get AID, including the parathyroid, pituitary and pancreatic alpha cells, for reasons that are not understood (Fig. 1a).

**Fig 1:**
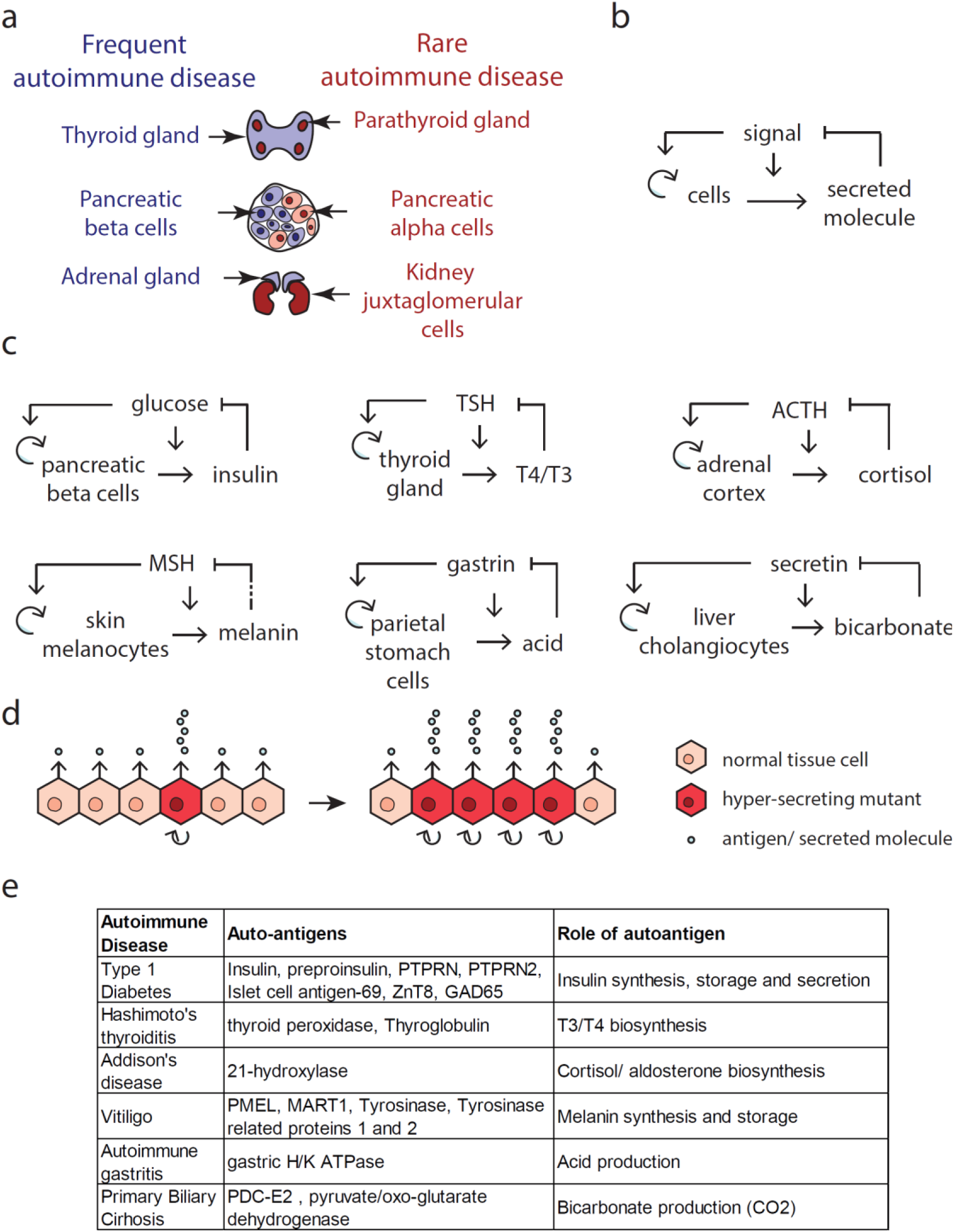
Regulatory circuits in secretory tissues that show prevalent autoimmune diseases link hyper-sensing of the input signal with hyper-secretion and hyper-proliferation. a) Some endocrine tissues are prone to autoimmune diseases, while others are not. b) The regulatory motif common to all tissues. c) The specific implementation of the motif in each tissue. Dashed line for melanin is an indirect affect in which melanin reduces UV that induces MSH. d) Due to this circuit, mutant cells that hyper-sense the signal (dark red cells) have a larger growth rate and hyper-secrete. Due to their growth advantage, these mutant cells can expand, eventually causing physiological hyper-secretion, and hence loss of homeostasis (hypo-signal). e) The autoimmune diseases related to the circuits in panel c have autoantigens in the biosynthesis and secretion pathways of their corresponding secreted factor. See Table S1 for details.

More generally, the fundamental origin of AID is currently unknown. One point of view is that the T-cells that attack specific organs are escapees of protection mechanisms that act to prevent autoimmunity. These protection mechanisms include central tolerance in the thymus and peripheral tolerance (1). Editing of self-reactive T-cells in the thymus not only removes T-cells with high affinity to self-antigens, it also promotes the development of FOXP3-dependent regulatory T-cells (Tregs) (2). These Tregs suppress the activity of self-reactive T-cells. Indeed, genetic predisposition to autoimmune disease lies primarily in genes related to Treg development and function and MHC class-II genes, as well as thymus function (1). Other protective mechanisms include immune checkpoints, as evidenced by the occurrence of auto-immune diseases in patients receiving immune checkpoint blockade therapy (3).

In addition to protective mechanisms, research has pointed to mechanistic process involved in promoting autoimmune disease. This includes post-translational modifications of auto-antigens. Examples include deamidation of gluten in celiac disease, pro-insulin in T1D, and citrullination of autoantigens in RA (4). Furthermore, misfolded proteins can be presented by MHC class II and thus seen as non-self by T-cells and not recognized by Tregs (5). Translational errors giving rise to defective ribosomal products (DRiPs) were also shown to act as self-epitopes in autoimmune disease (6). Certain autoantigens may be more likely to be presented by certain MHC class I molecules, and more susceptible to environmental PTM.

Another risk factor is viral infection. One theory, called molecular mimicry (7), is that pathogens have proteins that resemble certain self-proteins, setting off autoimmune attack. A related hypothesis is that viral infection weakens immune tolerance mechanisms and sets off an autoimmune response.

A different point of view is that the self-reactive T-cells play an important physiological role (8–12). This is supported by evidence that self-reactive T-cells are found in healthy individuals, including T-cells that recognize beta cells, melanocytes and thyroid cells (13–17). The young age of onset and devastating nature of organ-specific autoimmune diseases carries a large evolutionary cost. Thus, their relatively high prevalence suggests that they originate from some process which is essential for human biology. What this process exactly is, however, is unknown. It would be important to identify an essential biological process that requires self-reactive T-cells towards specific antigens in specific endocrine glands. Such a process would supply the healthy counterpart to autoimmune diseases.

Here we propose such an essential process. We start from the point of view of tissue-level feedback loops. We argue that tissues that turn over and that secrete factors such as hormones must have feedback control of their cell proliferation, in order to maintain proper organ size and secretion rate (18). Such feedback inherently shows a fragility to mutant cells that mis-sense the feedback signal: such cells both hyper-secrete the hormone and begin to rapidly proliferate (19). The mutant proliferation poses a threat because once the hyper-secreting mutant clone reaches a large enough size, it can lead to physiological dysregulation.

We hypothesize that in order to prevent these mutants from expanding, each organ has a dedicated surveillance mechanism based on autoimmunity. In this system, self-reactive T-cells preferentially eliminate the hyper-secreting cells. This postulated ‘autoimmune surveillance of hyper-secretion mutants’ (ASHM) is essential to prevent mutant clone expansion, but makes the tissue prone to tissue-specific autoimmune disease. We test several predictions of this hypothesis. The first prediction is that the antigens are in key enzymes in the production of the secreted molecule. The second prediction is that endocrine tissues that rarely get autoimmune disease instead are prone to age-dependent mutant clone-expansion diseases with hyper-secretion. A prime example is the parathyroid gland which very rarely gets autoimmune disease, but frequently gets tumors that hyper-secrete parathyroid hormone, causing hyper-parathyrodism. We discuss experimental tests of this hypothesis and clinical implications.

## Results

### Secretory organs prone to autoimmune disease share a common feedback circuit motif

Autoimmune diseases are classified into systemic diseases such as lupus, rheumatoid arthritis and systemic sclerosis, and organ-specific diseases such as type-1 diabetes (T1D), Hashimoto’s thyroiditis and autoimmune Addison disease. Here we focus on endocrine and exocrine organ-specific diseases. We list organ-specific autoimmune diseases of high prevalence (more than 0.1% lifetime prevalence) in Table S1. In these diseases. T-cells attack pancreatic beta cells, thyroid, adrenal cortex, melanocytes, gastric parietal cells and liver cholangiocytes.

We consider these systems from the point of view of tissue-level regulatory circuits. All of these tissues have cell turnover, in which cells proliferate and are removed on a timescale of months. As shown by Karin et al (18), in order to maintain organ size, cell proliferation and removal must be precisely balanced, and hence must be controlled by a feedback loop. This feedback loop prevents exponential growth or collapse of the tissue. In order to provide organ functional mass that matches physiological needs, the feedback must operate based on input signals that are related to the tissue function.

We therefore sought for each organ system what signals control cell proliferation and hormone secretion. We find that these organs share a regulatory motif (Fig. 1b,c). The input signal (e.g. blood glucose in the case of beta cells) increases both the secretion of a hormone (e.g. insulin) and the growth rate of the secreting cells by increasing proliferation and/or reducing apoptosis. The secreted factor, in turn, regulates the input signal (e.g. insulin reduces blood glucose).

Thus, *secretion and proliferation are linked in these cells*. If there is too little hormone, the circuit makes the cells secrete more hormone, and also proliferate to enable larger hormone supply over longer timescales.

Analyzing these circuits, we note that they share a fragility to mis-sensing mutant cells (Fig. 1d). A mis-sensing mutant is a cell with a mutation in the sensing pathway that causes it to over-sense the signal. As a result, the mutant cell both hyper-proliferates and hyper-secretes. An example in beta cells are mutants in glucokinase, that cause beta cells to over-sense glucose and as a result to hyper-secrete insulin (20–22). In the thyroid, TSH-receptor mutants grow into a nodule that hyper-secretes thyroid hormone (23).

We use the word mutant, but the over-sensing might in principle be due not to a genetic change but instead to a long-lasting epigenetic aberration that is passed on to daughter cells.

Because mis-sensing mutants have a higher growth rate than other cells in the organ, they compete with other cells and can expand to form clones. When the clones reach a large enough fraction of the total cell population, their hypersecretion begins to interfere with homeostasis.

At first sight, it might seem that such mutations arise rarely. However, when considering the mutational target and mutation rates, it is easy to see that these mutations are practically unavoidable. To estimate the frequency in which such mutants arise, we need to multiply the mutation rate by the number of cells, and to estimate the cell proliferation rate. We illustrate this for the case of the thyroid. Mutation rate is about p=10^−9^/bp/division, and thyroid follicular cells number about N=10^10^. Thus, any given mutation will arise about ten times every thyroid turnover time. There are at least 45 different activating mutations in the TSH receptor that give rise to thyroid hypersecreting nodules (24).

Thyroid cells divide at a low rate in adults in normal physiological states, but can rapidly expand as a physiological adaptation, for example in the case of goiter formation due to insufficient iodine intake (25,26). The cells divide more rapidly in children, let’s say once per year (27). Taking a turnover time of one year means that every possible point mutation in the genome arises about ten times per year in the thyroid. A typical mis-sensing mutation such a mutation in the TSH receptor makes the thyroid cells proliferate at, say, twice the wild-type rate (e.g. divide twice per year) (28). The mutant population will thus grow a thousand-fold in 5 years and a million-fold in 10 years, so that a single mutant can take over the population in less than 15 years. Such mutant takeover will raise thyroid hormone to toxic levels which can be lethal (29), as in toxic thyroid nodules. Thus, mutant expansion in young individuals seems to be an almost unavoidable threat.

An even more fundamental situation in which mutant cells are a concern is embryonic development. For example, to produce the 10^9^ beta cells in the body, requires (at least) 10^9^ divisions, and thus mutant cells are likely to be present already at birth. Indeed, such somatic mutants result in focal hypoglycemic hyperinsulinism in neonates, causing expansion of hyper-secreting, hyper-proliferative beta cells, which results in potentially lethal hypoglycemia (30). Mutant expansion is more likely the larger the number of cells, as in large organs like the thyroid and adrenal. It is also more likely the faster the cell division rate, as in childhood and puberty and in conditions that increase proliferation such as iodine deficiency in the thyroid.

In cases where the fitness cost of hypersecretion is severe enough, there is scope for a powerful mutant-resistance system, even if that system itself carries fragilities.

### To resist homeostasis disruption, we propose a mechanism based on autoimmune surveillance of hyper-secreting mutants

Here we propose a mutant-resistance system based on autoimmunity, which we call *autoimmune surveillance of hypersecreting mutants*, ASHM. The idea is that the adaptive immune system detects and removes cells with high expression of proteins in the secretion pathway. In this way, autoimmune cells give a growth disadvantage to hyper-secreting cells relative to the other cells.

An ASHM mechanism makes the following predictions: i) the autoantigens must be in the pathway making the secreted molecule, ii) endocrine tissues with rare autoimmune disease (weak ASHM) should show common expansion of hyper-secreting mutants.

### Autoantigens in endocrine autoimmune diseases are often in the secretion pathway

To test the prediction that the autoantigens must be in the pathway making the secreted molecule, we listed the major antigens in the endocrine and exocrine autoimmune diseases in Fig. 1e (see also Table S1). All but one of the known antigens are in the secretion pathway.

In T1D, for example, the major autoantigens, preproinsulin and protein tyrosine phosphatase, are both in the insulin synthesis pathways. In Addison’s disease, the major autoantigen, 21-hydroxylase, is a rate limiting step in cortisol and aldosterone synthesis. In Hashimoto’s thyroditis, the autoantigens are thyroglobulin and TPO, both critical steps in thyroxine synthesis. In vitiligo, the autoantigen is the rate limiting step in melanin production. In autoimmune gastritis, the autoantigen is the acid transporter.

There is one exception in which the antigen is not known to be related to the biosynthesis of the secreted molecule, namely GAD in T1D. This case may require additional explanation, such as epitope spreading, as suggested by Codina-Busqueta et al (31).

These antigens are specific to their cell type. It is thus instructive to consider a case in which the auto-antigen is present in many different cell types: the case of primary biliary cirrhosis (PBC).

PBC is an autoimmune disease in which T-cells attack the liver cholangiocytes. The auto-antigen is a mitochondrial protein (anti-mitochondrial antigen, or AMA). It is a well-known puzzle that a mitochondrial antigen, present in all tissues, would lead to attack on specific cells, the liver cholangiocytes (32).

The present concept can explain this puzzle. Cholangiocytes produce bicarbonate for bile secretion (Fig. 1c). According to the ASHM hypothesis, the antigen should be in the bicarbonate secretion pathway. Indeed, the auto-antigen is a regulatory subunit of pyruvate-dehydrogenase and oxo-glutarate dehydrogenase. These dehydrogenases are the main producers of C0_2_ in the cell, through glycolysis and the TCA cycle. C0_2_ is the basis for production of bicarbonate. Thus, cholangiocyte mutant cells that hyper-secrete bicarbonate are likely to have more of this antigen than their neighbors. These mutant cells can be removed by ASHM, without affecting other tissues in which there is no growth-advantage for cells that upregulate this antigen. Thus, the ASHM hypothesis offers a functional explanation for the mitochondrial antigen in this disease.

### Endocrine tissues that rarely get autoimmune disease are prone to mutant-expansion diseases

The second major prediction of the ASHM hypothesis is that secretory organs that *lack* prevalent autoimmune disease, should be prone to *non-transient expansion of hyper-secreting mutant cells*. These organs will therefore show prevalent diseases of hypersecretion.

To test this, we consider endocrine systems that very rarely get auto-immune diseases and for which cell growth regulation is well-studied (Fig. 2a, Table S2). This includes the parathyroid gland, the pituitary gland, renin-secreting cells in the kidney and alpha-cells in the pancreas.

**Fig. 2:**
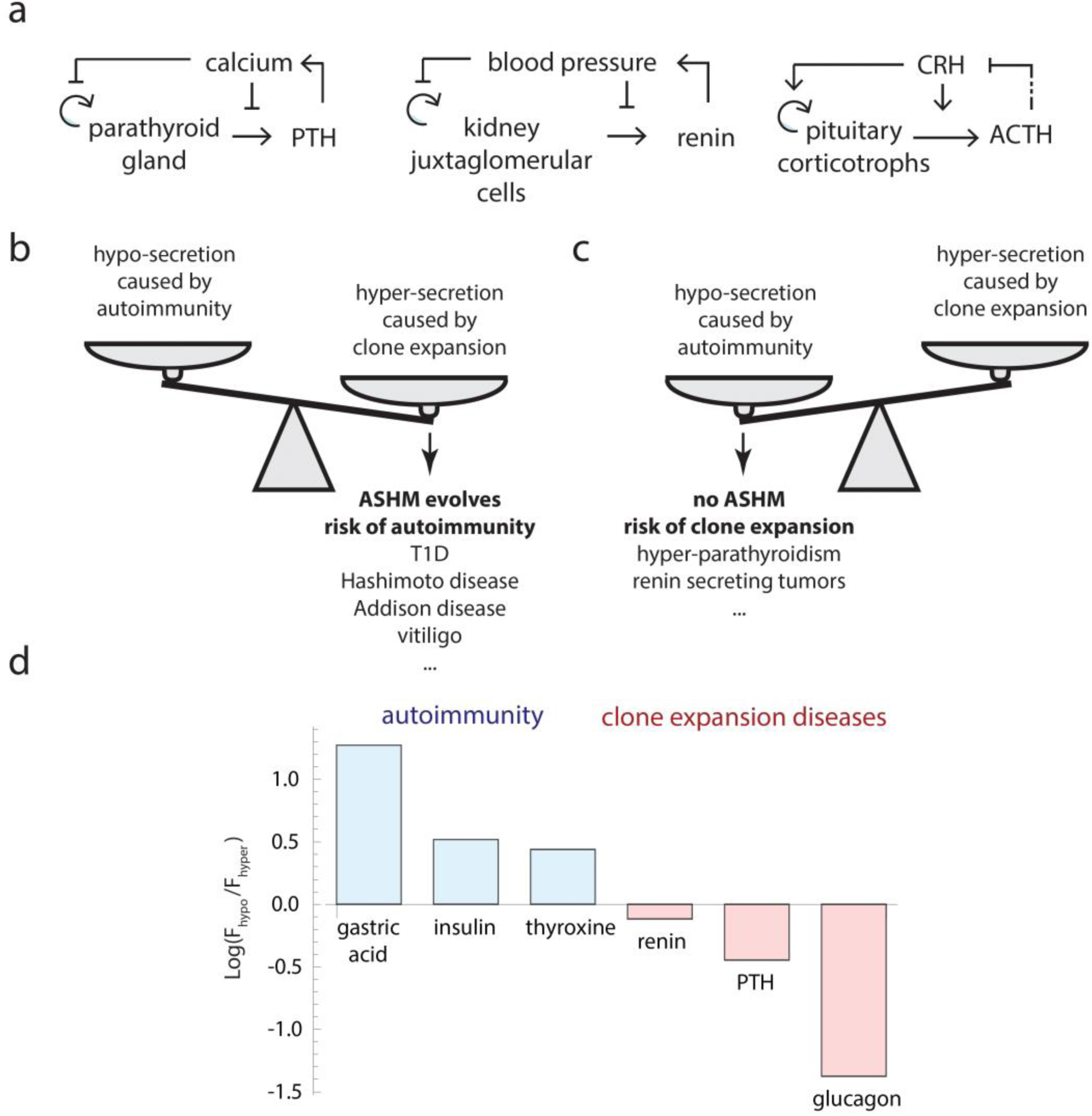
Circuits for three endocrine tissues that show autoimmune disease very rarely. a) Circuits for the parathyroid, renin and pituitary systems. Dashed line is the effect of ACTH inducing cortisol which inhibits CRH secretion. b) When the cost of hyper-secretion exceeds the cost of autoimmunity, ASHM can be selected. c) Otherwise, ASHM is not selected and diseases of hyper-secreting clone expansion are expected. d) A measure C for cost of hypersecretion versus equivalent hypo-secretion for six systems. Blue: systems with prevalent autoimmune disease, red: systems with rare autoimmune disease. For molecule x, C is F_hypo/F-hyper, where F_hypo (F-hyper) is the ratio of the normal concentration of x to the lethal level at low (high) perturbations. (Methods).

In some of these systems, the circuit is slightly different: the input-signal inhibits proliferation and secretion, instead of activating them as in Fig. 1. The circuit is still prone to mis-sensing mutant expansion. Such expansion causes hyper-regulation of the input signal, as opposed to hypo-regulation in the circuits of Fig. 1.

We hypothesize that these systems have rare autoimmune disease because they lack an ASHM mechanism, or show a weak version. This predicts that such tissues will show relatively frequent *expansion of mutant clones which hyper-secrete*.

Such expansion occurs frequently in the case of the parathyroid gland. Here, mutant cells that under-sense calcium proliferate, and cause hyper-calcemia, in a disease called primary hyperparathyroidism. This disease is prevalent at old age, in as many as 1/50 postmenopausal women (33).

Clone expansion disease also occurs in the renin system, called renin-secreting juxtaglomerular cell tumors, or reninomas. These tumors hyper-secrete renin and cause high blood pressure. Reninomas have a prevalence of about 0.01% (34).

The pituitary gland is another example, which gets autoimmune disease extremely rarely, with about 500 cases reported in 30 years (35). In contrast, mutant expansion is common in the pituitary, estimated at about 20% of the population (36,37). The chance per cell for an ACTH-secreting pituitary adenoma is about 250-fold greater than a cortisol-secreting adenoma in the much larger adrenal cortex (SI section 1). Such hypersecreting adenomas lead to endocrine problems: Cushing’s disease when ACTH is hyper-secreted (38), acromegaly when growth hormone is hyper-secreted (39), Hypogonadism when sex hormones are secreted and hyperthyroidism when TSH is secreted. The prevalence of such hypersecreting adenomas totals about 0.01%.

Thus, tissues with rare autoimmune disease seem to show prevalent mutant hyper-secretion diseases. We predict from this picture that a similar mutant-expansion disease might occur in alpha cells (whose regulatory circuit is more complicated, SI section 2), causing a phenotype of hyper-glucagonism and hyper-glycemia. Such expansion phenomena have been reported, although other explanations have been offered (40–42).

One may speculate about a principle that dictates which tissues get autoimmunity versus which get clone-expansion diseases: If the fitness cost of hyper-secretion or hyper-expansion is higher than the fitness cost of autoimmunity, ASHM will evolve (Fig. 2b) (43). In this case, ASHM will prevent mutant takeover in many individuals, at the cost of young-onset autoimmune diseases in a few. For example, the fitness cost of hyper-insulinemia is very large because hypo-glycaemia is lethal. ASHM prevents this lethality, at the cost of T1D which causes lethality to a small fraction of the population.

Conversely, if the fitness cost of hyper-secretion/hyper-expansion is lower than the fitness cost of autoimmunity, ASHM will not evolve (Fig. 2c). These systems will instead exhibit mutant-takeover diseases of hyper-secretion, whose prevalence rises with age. For example, the fitness cost of alpha-cell mutant takeover with hyper-glucagonemia seems to be low, because hyper-glycemia can be tolerated or corrected by insulin. Hence no ASHM evolves against alpha-cells. Similarly, hyperparathyroidism is tolerable (mild hyper-calcemia leading to gradual bone damage and cognitive effects), as compared with acutely lethal hypo-calcemia that would result from an auto-immune disease.

To test this principle requires knowing the fitness costs of hyper- and hypo-secretion, which are difficult to estimate. As a first approximation, we sought a proxy for the relative fitness costs by estimating the deviation from normal ranges in which lethality occurs. For example, lethal hypocalcemia occurs when free calcium drops by a factor of about F_hypo=1.1 (normal range is 1.1-1.3 mM and lethality occurs below 0.9 mM). Lethal hypercalcemia occurs when F_hyper=2.5-3 (above 2.5-3 mM). No ASHM is predicted in this case because F_hypo is much smaller than F_hyper.

Conversely, we reasoned that ASHM would be favored if F_hyper is smaller than F_hypo, because mutant-expansion leading to hyper-secretion would be dangerous compared to a similar level of hypo-secretion caused by autoimmune surveillance. For insulin control of glucose, for example, lethality occurs below 3 mM blood glucose, which is 1.7-fold below the normal 5 mM, whereas a clinical high level (risk of ketoacidosis) is estimated at 2.8-fold above normal.

We found estimates for six of the relevant factors in this study (Fig. 2d, Methods). For gastric acid, insulin (glucose) and thyroxin, F_hyper is smaller than F-hypo, consistent with autoimmune disease. In contrast, for calcium (PTH), blood pressure (renin) and glucagon, the opposite is true, consistent with mutant-expansion diseases. These rudimentary estimates support the proposal that AID is likely in tissues where hypersecretion carries a high fitness cost, whereas mutant-expansion diseases is likely in tissues where hypo-secretion carries a higher cost.

Additional factors may include the probability for a mis-sensing mutant, which is smaller in small tissues of under 0.1g such as pituitary and parathyroid (and hence less need for ASHM), compared to large tissues on the order of 10g such as the thyroid and adrenal cortex (and hence more need for ASHM).

The occurrence of AID in these organs in immune checkpoint therapy or in congenital mutations (e.g. in AIRE) leading to polyglandular autoimmune disease suggests that these organs may have a reduced amount of ASHM rather than no ASHM at all. The potential mechanisms for certain organs to have reduced ASHM include at least two possibilities: greater expression of the relevant tissue protein in the thymus under AIRE control, trimming potentially reactive T cells from the repertoire. Alternatively, the relevant proteins could lack sequences likely to bind to host MHC molecules, or the organs could have reduced MHC class I pathways. These possibilities are experimentally testable.

### ASHM requires cytotoxic T-cells to differentially remove cells that make more auto-antigens then their neighbors

We now discuss the plausibility of ASHM in terms of what T-cells can do. ASHM requires the immune effector cells to differentially recognize and remove cells that make more auto-antigens then their neighbors.

Such differential killing is unlikely if the antigen recognition is very strong. For strong antigens, such as certain foreign antigens, a single or a few pMHC per cell have been shown to activate CD8 T cells (44,45).

However, for weaker antigen recognition, there is evidence for differential sensitivity based on the number of MHC molecules presenting the antigen on the cell surface. A cell tracking experiment by Halle et al showed cooperativity, in which cell killing is a steep function of the number of T-cell contacts with the target cell with Hill coefficients exceeding 7 (46,47). One basis for such cooperativity is evidence that that T-cell receptors (TCRs) work cooperatively in the same T-cell, so that TCR activity spreads in a TCR nanocluster (48).

Indeed, T-cells with very strong autoantigen binding are trimmed from the emerging repertoire in the thymus, based on low-level expression of peripheral organ-specific proteins in thymic stomal cells under control of AIRE. This is consistent with the notion that the TCR repertoire is adjusted for recognition selectivity where a quantitative difference in protein expression of 1-2 fold would provide sufficient discrimination to achieve the postulated detection of hypersecretors.

We conclude that T-cells may be capable of differential killing according to auto-antigen levels required for ASHM, especially if the recognition is moderate.

### The public T-cell repertoire includes TCRs that recognize auto-antigens involved in endocrine autoimmune disease

Since ASHM is predicted to be common to all individuals, a further prediction is that TCRs that are responsible for autoimmune diseases and are expanded in individuals with these diseases, should also be present in all healthy individuals, namely in the public T-cell repertoire. To test this, we sought TCR sequences derived from patients or mouse models of the diseases, and which interact with the auto-antigens. Such TCR beta-chain sequences were identified for T1D (proinsulin, ZNT8, GAD65 and islet autoreactive TCRs) (13,49), Hashimoto’s thyroiditis (thyroglobulin) (50) and vitiligo (PMEL, MART1) (51,52).

We compared these sequences to databases of public T cell receptor sequences that are shared in healthy humans or healthy mice ((53), Table S3). We find 30 perfect matches for TCRs from T1D, Hashimoto’s and vitiligo (Methods). Strikingly, four of these perfect matches occur in both the mouse and human datasets, in a set of 92 TCR sequences common to both organisms. We conclude that the TCRs that participate in the disease process are also found in the public TCR repertoire, consistent with their postulated physiological role in ASHM.

### Mathematical modelling suggests that ASHM can eliminate mutant cells while killing few normal cells

Finally, we turn to mathematical modelling in order to ask what features of ASHM are essential in order to protect from hypersecreting mutants. The conclusions of this analysis is that ASHM can protect against mis-sensing mutants even when it kills at a rate that is smaller than the natural turnover of the tissue. For ASHM to work effectively, it needs to attack cells at a rate that is strongly nonlinear in antigen level (cooperative response). However, too high killing rates leads to tissue loss and autoimmunity, illustrating the need for tight regulation of ASHM.

To model ASHM, we consider the growth-rate of a cell clone which senses the input signal *s* (Methods). Mutant cells sense a distorted input signal, which we call the ‘perceived signal’, so that they sense *u* times more signal than wild-type cells. Cells have a growth rate and secretion rate that depends on the perceived signal. Cells are also removed by ASHM, which kills cells at a rate which is a function of their antigen level, which in turn is proportional to their secretion rate (Fig. 3a-b). ASHM killing is cooperative as described by a Hill function with coefficient *n*. As mentioned above, experiments suggest high cooperativity. The baseline killing rate of the ASHM system is described by the parameter *γ*, the rate of cell deaths caused by surveillance relative to cell death by natural turnover.

**Figure 3:**
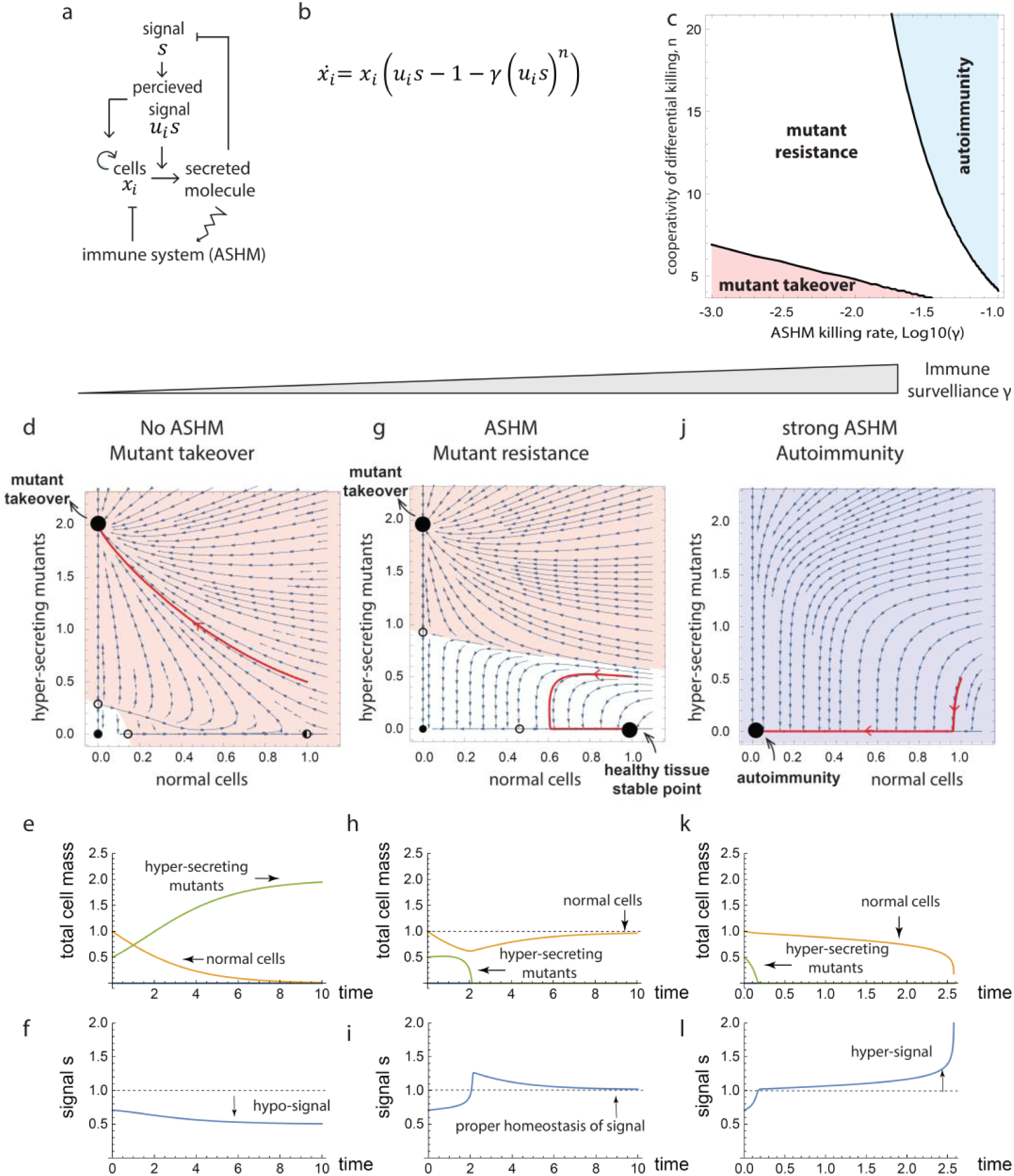
Mathematical model indicates that ASHM can protect against mis-sensing mutants with cooperative differential killing that is infrequent compared to natural turnover. a) Scheme for ASHM mechanism. b) Equation for ASHM model, in which clones of cells *x*_*i*_ with mis-sensing distortion *u*_*i*_ are subject to ASHM with killing rate *γ* and cooperativity *n* (Methods). c) Mutant takeover (*u* = 3) occurs when ASHM is not cooperative (low *n*), autoimmunity (killing of >50% of cells) occurs when ASHM is too strong (high *γ*). A wide parameter range (white) allows for removal of mis-sensing mutants without sizable loss of tissue. Phase plane analysis shows mutant and wild-type populations (*u* = 3, *u* = 1 respecitevely) for: (d,e,f) low *γ* (*γ* = 10^−4^) showing mutant takeover and hypo-regulation of signal, (g,h,i) *γ* = 0.01 showing mutant elimination and normo-regulation unless mutant population is large, and j,k,l) strong ASHM (*γ* = 0.1) showing loss of tissue and hyper-regulation of signal. (Methods).

We asked whether such an ASHM mechanism can remove all possible mis-sensing mutants (all values of the perceived-signal parameter *u*). In other words, we asked whether ASHM can provide an ‘evolutionary stable strategy’ in which no mis-sensing mutant can invade and expand in a wild-type population of cells (Methods). We find that ASHM can indeed remove all possible mis-sensing mutants provided that the ASHM killing rate *γ* equals 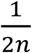, where *n* is the Hill cooperativity of immune discrimination. Given the high observed cooperativity of cytotoxic T-cells, n>7, the ASHM can work with a small killing rate relative to the natural cell removal rate in the tissue (*γ* < 0.1).

To illustrate the dynamics of ASHM, in Figure 3 we plot the dynamics of a hyper-sensing mutant clone that senses twice the true signal (*u=2*) (Methods). Without ASHM, a single mutant cell in a wild-type tissue expands exponentially and takes over (Fig. 3d,e). As a result, homeostasis is lost (Fig. 3f). The same mutant rapidly vanishes with ASHM (Fig. 3h,i). If the mutant population reaches a critical size, however, ASHM can no longer control it, and the mutant takes over the tissue, as shown in a phase portrait in Fig. 3g (orange region). Thus, ASHM is a frequency-dependent mechanism: if all cells are mutant, ASHM cannot distinguish between them and selectively eliminate mutants. Thus, ASHM cannot help in cases of germline mis-sensing mutations.

ASHM can also eliminate multiple simultaneous mutant clones for plausible parameters in the range of low killing *γ* = 0.05, and high cooperativity *n* = 10 (Fig. 3c, Methods). One can prove that ASHM works for a general class of feedback models (Methods).

Autoimmune disease can be modelled by raising the killing rate *γ*. This represents a situation where B-cells and memory T-cells are activated leading to sustained immune attack and memory.

## Discussion

We presented the hypothesis that endocrine organ-specific autoimmune diseases originate from an essential physiological mechanism that removes deleterious hyper-secreting mutants. We find that endocrine and exocrine glands prone to autoimmune disease share a regulatory motif in which the signal that they control increases both cell proliferation and secretion. Such feedback gives mutant cells that mis-sense the signal a growth advantage, leading to clone expansion that can ruin homeostasis. To remove these mutants in cases where signal perturbation is life-threatening, we propose an autoimmune surveillance of hyper-secreting mutants (ASHM) mechanism (Fig. 4). We find evidence for the basic requirements of this mechanism: the major autoantigens in the autoimmune disease are from proteins in the secretion pathways, T-cells against these autoantigens are in the public T-cell repertoire shared by all healthy individuals, and immune killing can differentiate hypersecreting cells from other cells. This picture predicts that endocrine tissues that rarely get autoimmune disease are instead prone to hyper-secreting mutant expansion, explaining the prevalence of hyperparathyroidism, hyper-secreting pituitary adenomas and several other hormone-secreting tumors. We show mathematically that ASHM can protect against all possible mis-sensing mutants (that is, mutant cells that perceive a distorted input signal) with a relatively low killing rate compared to the natural tissue turnover.

**Figure 4:**
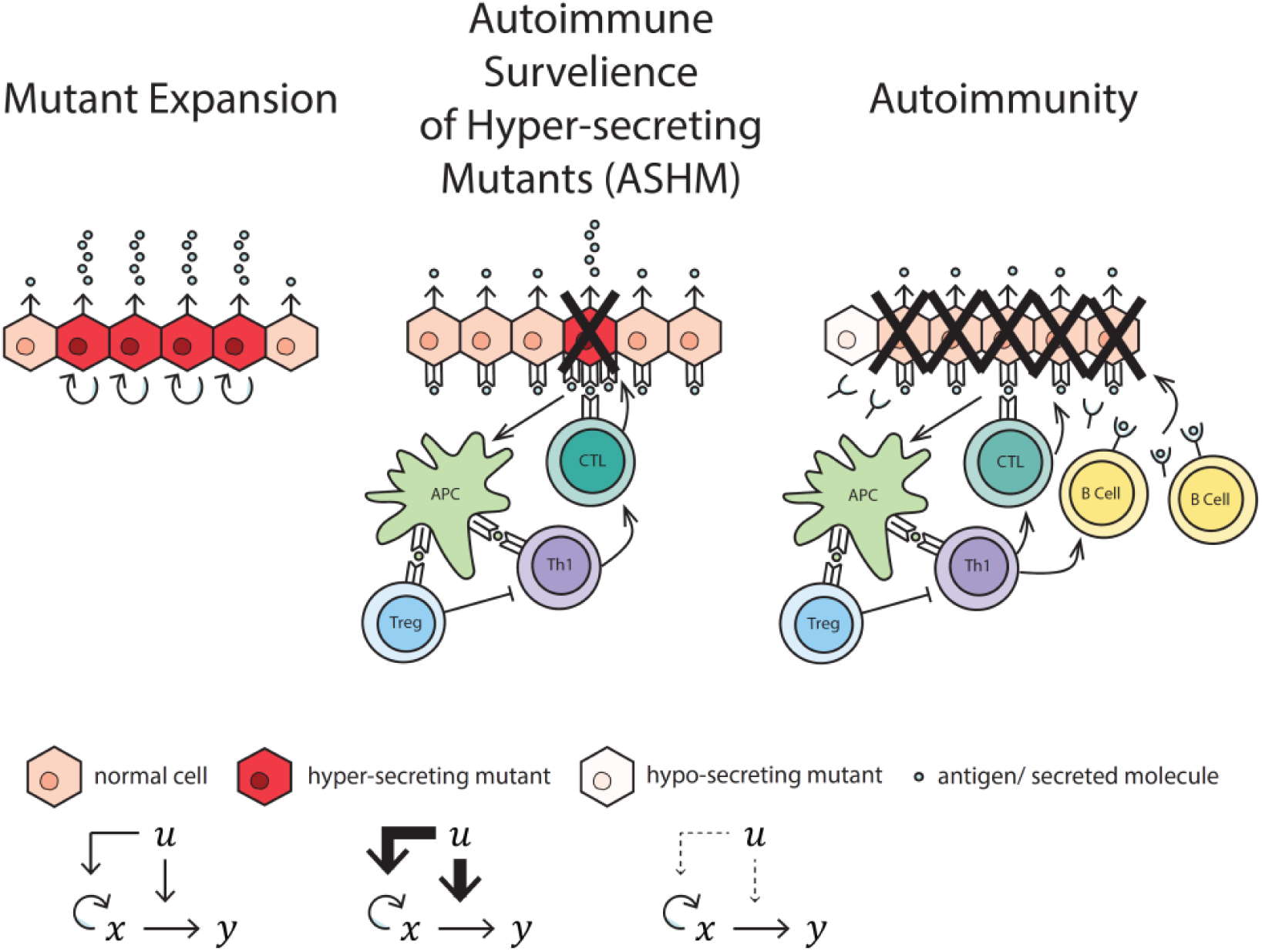
The ASHM mechanism (autoimmune surveillance of hypersecreting cells) can prevent hypersecreting clone expansion, at the cost of risk of autoimmune disease. ASHM T cells recognize antigens in the secretion pathway to selectively eliminate cells that secrete more than their neighbors. When over-activated, the same cells can set off a persistent immune response including B-cell activation that kills much of the tissue, except for hypo-secreting clones.

Together with its essential mutant-removal function, ASHM poses a risk because it can degrade into autoimmune disease. Onset of autoimmune disease can occur, for example, if there is a change in the tissue that increases antigen presentation (54–56). Possibilities include tissue damage that causes release of antigen, cell damage that leads to senescence (57), or cell overload that leads to expression of strongly immunogenic variants of the antigen (6,9). These changes might lead to B-cell activation and response as if the autoantigen had a microbial origin. The danger for this might increase if a viral infection coincides with the tissue damage.

ASHM is presumably tightly regulated by Tregs and immune checkpoints, and held at an intermediate activity that avoids killing too many healthy cells and at the same time eliminates hypersecretors. Thus, weakening immune checkpoints as in cancer immune-therapy commonly leads to endocrine autoimmune diseases (3). Similarly, mutations in genes involved in immune checkpoints or Treg development and function are known to increase the risk for autoimmunity (1).

In addition to ASHM, tissues with the present regulatory motif often have another mechanism against mis-sensing mutants. This mechanism is cell-autonomous biphasic control, in which the signal promotes growth of cells at low levels but kills cells at very high signal levels (19). For example, glucose drives the growth of beta cells at low to medium concentrations, but drives their death through glucotoxicity at high concentrations. Similarly, calcium has a biphasic effect on the parathyroid cells that control it. This biphasic mechanism is useful to prevent extreme hyper-sensing mutants: these mutants perceive high levels of signal and kill themselves. However, biphasic control is vulnerable to mild mutants, which mis-sense the signal at below its toxic levels; removing such mild mutants require an additional mechanism such as ASHM. This observation is in line with the mild nature of the mis-sensing mutants found in the expanding clones of hyperparathyroidism (58). Biphasic control may not be feasible in tissues which need to sense the signal over a wide dynamic range, such as the TSH signals to the thyroid which can span three orders of concentrations (59,60).

The ASHM model predicts that self-reactive T-cells against the specific autoimmune antigens will be found in healthy people. Several studies support this prediction (14–17). For example, Culina et al finds fully functional cytotoxic islet-reactive CD8+T cells in the blood of healthy people, in amounts similar to people with T1D (in T1D there are more of these cells in the pancreas compared to in healthy people) (13). Here we further find that T-cell receptor beta-chains against specific disease antigens are found in the public T cell repertoire shared by healthy individuals. Such self-reactive T-cells may be at play in the endocrine autoimmune diseases which are a common side effect of immune checkpoint therapy for cancer (3).

The ASHM mechanism is in line with the view that self-reactive immune cells play a role in tissue homeostasis and body maintenance (8–11). It proposes that self-reactive T-cells have an essential role in removing hyper-secreting mutants, and are not merely unwanted escapees of peripheral clonal deletion/anergy or due to errors of elimination in the thymus. Along these lines, another role of the proposed ASHM mechanism could be to trim away cells that hyper-secrete for non-genomic reasons, such as stochastic fluctuations in receptor number or other parts of the sensory or secretory apparatus. Such a trimming role is especially relevant in tissues with slow turnover, such as neurons, in which hyper-secreting cells could have a deleterious local effect for prolonged periods.

The ASHM mechanism can be experimentally tested. A critical test would be to examine the consequences of a systemic lack of T-cells, which is predicted to enhance the prevalence of hypersecreting mutant expansion in endocrine organs. ASHM can be further tested by seeking, in healthy organisms, autoreactive T-cells against the secretion pathway proteins among tissue-resident T-cells of the target organ or in adjacent lymph nodes. These T-cells should be active and kill cells in the healthy tissue. The killed cells should be preferentially those with high secretion. These experiments are feasible in principle, and can be carried out in the different tissues discussed here including thyroid, pancreas, adrenal cortex, skin and stomach. A relative lack of such ASHM tissue-resident T-cells is predicted in tissues such as the parathyroid, pituitary and the renin-secreting cells of the kidney. Identifying the molecular pathways for ASHM, and the nature of the participating T-cells, may provide targets to tune its strength (the parameter *γ*) by pharmacological means, allowing control of the tradeoff between mutant expansion and autoimmune disease.

## Supplementary Tables

**Table S1:** Endocrine autoimmune diseases.

| Autoimmune Disease | Cells | Prevalence | Input | Secreted factor | Auto-antigens | Role of autoantigen |
| --- | --- | --- | --- | --- | --- | --- |
| Type 1 Diabetes | Pancreatic beta cells | 0.1-1% | glucose | Insulin | Insulin | Insulin |
|  |  |  |  |  | preproinsulin (PPI) | Insulin precursor |
|  |  |  |  |  | Protein Tyrosine Phosphatase, Receptor Type N (PTPRN) | Insulin storage and secretion (61,62) |
|  |  |  |  |  | Protein Tyrosine Phosphatase, Receptor Type N2 (PTPRN2) | Insulin secretion regulation (63,64) |
|  |  |  |  |  | Islet cell antigen-69 (ICA69) | transport of insulin secretory granules (65) |
|  |  |  |  |  | zinc transporter 8 (ZnT8) | Insulin production, storage, and secretion (66,67) |
|  |  |  |  |  | Glutamic decarboxylase 65 (GAD65) | No known role in insulin secretion |
| Thyroiditis (Hashimoto disease) | Thyroid gland cells (thyrocytes) | 0.1-1% | TSH | T4/T3 | thyroid peroxidase (TPO) | T3/T4 biosynthesis (68) |
|  |  |  |  |  | Thyroglobulin (Tg) | T3/T4 precursor |
| Addison Disease | Adrenal cortex cells | 0.01 % | ACTH | Cortisol/aldosterone | 21-hydroxylase (CYP21A2) | Cortisol/aldosterone biosynthesis |
| Vitligo | Melanocytes | 0.1-1% | MSH | melanin | Melanocyte protein PMEL | Melanin synthesis and storage (69) |
|  |  |  |  |  | Protein melan-A (MART1) |  |
|  |  |  |  |  | Tyrosinase (TYR) | Melanin synthesis (70) |
|  |  |  |  |  | Tyrosinase related proteins 1 and 2 |  |
| Autoimmune gastritis | stomach parietal cells | 0.1-1% | Gastrin | acid | gastric H/K ATPase | Acid production (71) |
| Primary Biliary Cirrhosis | Cholnagiocytes |  | Secretin | Bicarbona te | PDC-E2 ,E3BP components of pyruvate/oxo-glutarate dehydrogenase | Bicarbonate production (CO2) |

Prevalence calculations are based on data from (71–75)
Autoantigens are from: T1D: (76) – T cell antigens. Hashimoto thyroiditis: (77) – autoantibodies with evidence for T cell reactivity. Addison Disease: (78) – autoantibodies. Vitligo (79) – T cell antigens. Autoimmune gastritis: (71), T cells antigen. PBC: (32) – autoantibodies.

**Table S2:** Tissues that rarely get autoimmune diseases.

| Cells | Input | Secreted factor | Autoimmune Disease | Prevalence | Hypersecreting Mutant expansion disease | Prevalence |
| --- | --- | --- | --- | --- | --- | --- |
| Parathyroid gland cells | Calcium | PTH | Autoimmune hypoparathyroidism | $10^{-3}$-$10^{-4}\%$<br>(Mostly as a part of autoimmune polyendocrine syndrome type 1) | Primary hyperparathyroidism | 0.1-1% (80) |
| Alpha cells | Insulin | Glucagon | No known disease | - | Hyperglucagonemia | ? |
| Kidney juxtaglomerular cells | Blood pressure | renin | No known disease | - | Renin-secreting renal juxtaglomerular cell tumor | 0.06% (34) |
| Corticotrophs | CRH | ACTH | Autoimmune hypophysitis | 0.0005% (81) | pituitary ACTH secreting tumors (Cushing) | 0.003% (38) |
| somatotrophs | GHRH | GH | Autoimmune hypophysitis | 0.0005% (81) | pituitary GH secreting tumors (Acromegaly) | 0.006% (39) |

**Table S3:**
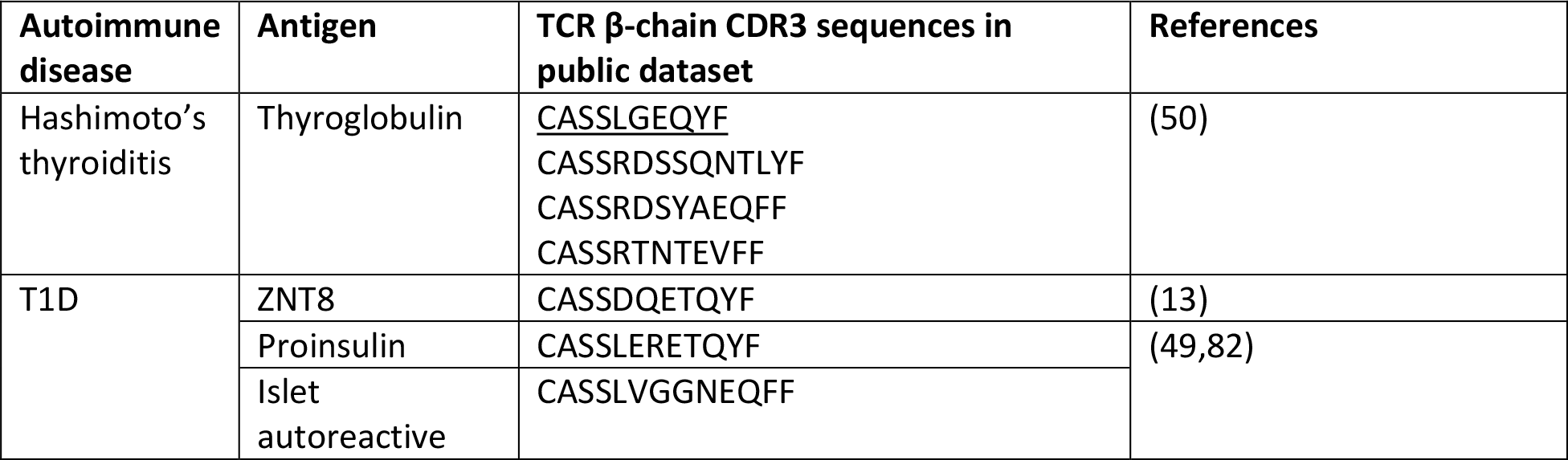
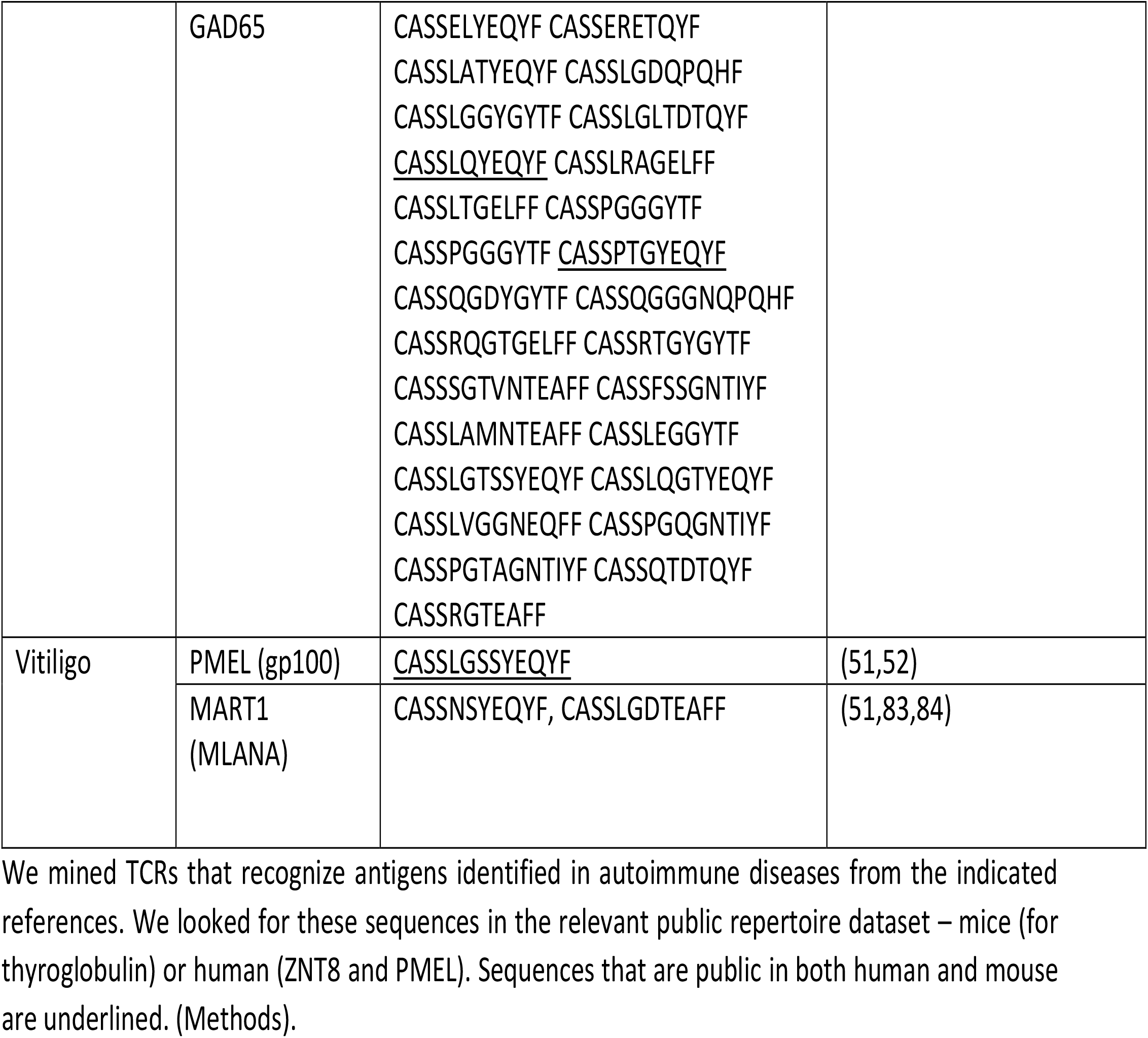
Autoimmune TCR sequences in public TCR repertoire databases.

## Methods

### Public TCR analysis

TCR beta chain sequences of T-cells experimentally found to react with disease autoantigens were retrieved by literature search (13,49–53,82–84). A total of 1010 sequences was thus collected. The public TCR beta chains were obtained from public-TCR databases as follows. A mouse database was retrieved from (14), using TCR sequences present in at least 25 of 28 mice studied (1006 sequences). A human public repertoire database, was built using the ADAPTIVE immuneACCESS database, using sequences shared by at least 400 people out of 785 (5016 sequences). We sought perfect sequence matches to the public TCR databases (perfect matches are listed in table 3). There were 92 sequences that appear in both mice and human public repertoire datasets. Perfect matches found in this restricted list are underlined in Table 3.

A statistical analysis for the likelihood of such perfect matches is not relevant for the present study, because the biological question is whether self-reactive TCRs are present in healthy individuals.

### Mathematical model for ASHM

Consider a cell clone *x*_*i*_ that has a net growth rate, proliferation minus removal, that rises with the perceived signal *u*_*i*_*s, f*(*u*_*i*_*s*), and a secretion rate that also rises with the perceived signal *g*(*u*_*i*_*s*) (Fig. 3a). ASHM removes cells at a rate that rises with antigen level, which is proportional to the cells secretion rate *g*. Hence, ASHM removal goes as *h*(*g*(*u*_*i*_*s*)). Thus,

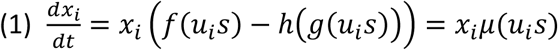

### Conditions for ASHM mutant removal

To find conditions in which ASHM can remove any mis-sensing mutant (any *u*_*i*_) on a background of wild-type cells (*u*_*i*_ = 1), we assume that signal is at its homeostatic set-point *s*\*, and require that the wild-type growth rate *μ*(*s*\*) is larger than that of any mutant *μ*(*u s*\*). Thus 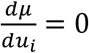 at *u* = 1, (and 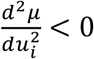, SI) providing the condition

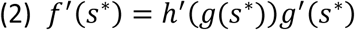

For linearized secretion and growth functions, *g* = *g*_0_*u*_*i*_*s, f* = *μ*_0_(*u*_*i*_ *s* − *s*_0_), and Hill-like killing with cooperativity *n, h* = *a*(*g*(*su*))^*n*^. The condition (2) becomes, for large n, approximately *γ* = 1/2*n*, where *γ* is the “ASHM strength” parameter, defined as the removal rate of wild-type cells by ASHM in units of their natural removal rate, 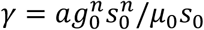. Thus, for the observed large T-cell cooperativity n∼10, the ASHM strength is small, 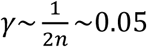. This means that ASHM can kill at about 5% of the natural removal rate and still be effective at removing mutants with any value of u.

To model the effect of mutant expansion on homeostasis, we use the model of Karin et al (18), namely that the signal is produced at rate *p* and removed by the secreted molecule *y* at rate *r*: 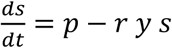, with *y* secreted by the different clones *dy*/*dt* = *b* ∑ *x*_*i*_ *g*(*u*_*i*_ *s*) − *α y*. For linear secretion and growth functions and Hill-like ASHM killing, the wild-type set point is 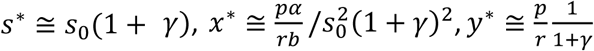. Thus the ASHM strength *γ* affects the set-point of the signal and cells. In autoimmunity (large *γ*), tissue is lost and the signal loses homeostasis.

### Simulation equations and parameters

We simulated the model equations using linearized growth secretion and Hill-like ASHM killing. We used a quasi-steady-state assumption for the dy/dt and ds/dt equations. We rescaled the parameters 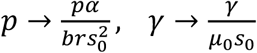, cell concentrations were normalized to the steady-state cell concentration, and time to tissue turnover rates, *t* → *μ*_0_ *s*_0_*t*. The resulting equation is 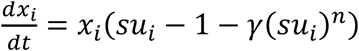 where 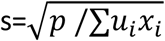. In Fig 4c, we defined the autoimmune disease region as the range of *γ*, n where this equation has no steady state solution except for zero cells. The mutant expansion region was defined as the range of *γ*,n in which a single mutant cell can invade the wild-type (*u* = 1) steady state (i.e. the steady state becomes an unstable saddle). We used a mutant with *u* = 3. In Fig 4d-n, we simulated a competition between a wild-type and a mutant population (*u* = 1 versus *u* = 3). The mutant initial concentration was half that of the wild-type cells, and parameters were *γ* = 10^−4^, 10^−3^, 10^−1^ and *n* = 10. All simulations were done using Wolfram Mathematica 11.3.

### Estimates of the fitness costs of hyper- and hypo-secretion

In order to estimate the fitness cost of hyper/hypo secretion, we use as a proxy the physiological concentration range of each secreted factor. We tabulated three parameters: 1. The normal concentration level, 2. The lowest concentration under which there is a severe and acute life danger 3. The highest concentration above which there is a severe and acute life danger. We didn’t consider long-term harmful effects of chronic unbalanced factor levels, because we reasoned that acute lethality carries a higher fitness cost than morbidity at old ages.

As a first approximation for the severity of the effect of mild hyper/hypo secretion respectively, we computed the ratios between the highest and normal concentrations as *F*_*hyper*_, and the normal to lowest concentrations *F*_*hypo*_, repsectively. In Fig. 2d we show log(*F*_*hypo*_/*F*_*hyper*_). Positive values indicate that the system is more sensitive to hypo-secretion than to hyper-secretion.

We found estimates for six of the relevant factors in this study. The values we use are:

#### PTH

we use ionized calcium level as a proxy. Normal levels are around 1.2 mM. severe hypocalcemia occurs below 0.9 mM (85). Hypercalcemic crisis occurs above 2.5 mM (86).

#### Insulin

we use glucose levels as a proxy. Normal glucose concentration is 5 Mm, while lethal low concentration leading to neuroglycopenic symptoms is 3 mM (87). High levels of insulin can lead to ketoacidosis, characterized by glucose levels higher than 14 mM (88).

#### Glucagon

we use glucose levels as a proxy, with the same normal and lethal-low concentrations as for insulin. However, the lethal effect of high glucagon is not ketoacidosis but hyperglycemic hyperosmolar syndrome, which happens above 33 mM glucose.

#### Renin

we use systolic blood pressure as a proxy. Normal level is 120 mmHg. Hypotension occurs below 90 mmHg and can be lethal because of fainting or shock. Hypertensive crisis can occur above 180 mmHg (values from the US National Heart, Lung, And Blood Institute). Considering diastolic blood pressure instead gives the same results.

#### Thyroxine

we use free T4 concentrations. Normal concentration is around 1.3 ng/dl. Lethal high value is 10.9 ng/dl (Thyroid storm, (89)). Lethal low levels occur around 0.1 ng/dl (danger of myxedema coma, estimation based on case reports, e.g. (90), and physiological distribution of FT4 levels in (91)).

For gastric acid, we use stomach pH levels. Normal baseline level is around 1.4 (92). We took low lethal level to be beneath pH of 1, which can lead to a potentially life threatening peptic ulcer (93). We could not find values for high lethal levels, for a conservative estimate we took the highest physiological level, pH 7, which can occur after a meal (92).

## Supplementary information

### S1: The probability of hypersecreting adenomas per cell is much higher in the pituitary than in the adrenal gland

The ASHM hypothesis predicts a tradeoff between organ-specific autoimmune disease (AID) and hyper-secreting mutant expansion diseases. One example is the case of pituitary gland corticotrophs that secrete ACTH vs. the adrenal cortex cells that secrete cortisol. Hyper-secreting mutants in both organs result in adenomas that cause a similar syndrome – pathologically high cortisol levels called Cushing’s syndrome.

ACTH-hyper-secreting adenomas in the pituitary gland, which is less prone to AID, are much more common per cell than hypersecreting adenoms in the adrenal cortex. To see this, note that the pituitary gland weighs ∼0.5g, and corticotrophs make up 15-20% of the anterior pituitary cells (Amar and Weiss, 2003). The two adrenal glands weigh ∼10g together of which ∼50% are cortisol-secreting cells (Neville and O’Hare, 2012; Wallig et al., 2017). Thus there are about 50 times more of the relevant cells in the adrenal. However, about 80% of Cushing’s syndrome cases are caused by pituitary tumors, and only 20% are from adrenal tumors (38). This means that per cell, the chance for a hyper-secreting tumor of pituitary corticotrophs is about 250 times larger than in adrenal cortisol secreting cells. This agrees with the predicted tradeoff.

### S2: Regulatory circuits for cell secretion and growth

In this section we provide details about the tissue feedback circuits that appear in Fig. 1c and 2a.

#### Beta cells

Beta cells secrete insulin, the main regulator of blood glucose. Insulin acts to reduce blood glucose by inducing uptake by peripheral tissue and reducing glucose production by the liver. An increase in blood glucose concentration stimulates the secretion of insulin by pancreatic beta cells. In addition, glucose increases beta-cell growth in rodents and humans (94,95). At very high levels, glucose causes beta cell functional decline and death, in a process called glucotoxicity. This biphasic effect of glucose can act to protect against strong mis-sensing mutants (Karin 2017).

#### Thyroid gland

Thyroid gland follicular cells secrete T4 and T3 in response to thyroid stimulating hormone (TSH) secreted by the pituitary thyrotropes. T4 is converted into T3 in peripheral tissue cells, controlling many aspects of metabolism. T4 and T3 in turn suppress TSH secretion, forming a negative feedback that keeps thyroid hormone concentration at homeostasis (96). TSH is also the major growth factor for thyroid gland cells (25,97). Thyroid growth provides a long-term compensation mechanism that can result in thyroid goiter formation, for example, as in the case of iodine deficiency.

#### Adrenal gland

The adrenal cortex cells found in the zona fasciculate secrete cortisol in response to ACTH stimulation, secreted by the pituitary corticotrophs. Cortisol suppresses ACTH secretion creating a negative feedback loop which regulates stress response (98). ACTH is the main growth factor of adrenal cortex cells (99).

#### Skin melanocytes

Skin melanocytes produce melanin, a pigment that protects against UV radiation, and transfer the pigment granules to surrounding keratinocytes. This process is regulated by melanocortins, including melanocyte stimulating hormone (MSH), secreted by the pituitary gland and skin, and ACTH (100). The negative feedback in this circuit is indirect: UVB radiation increases MSH receptor activity and MSH/ACTH precursor production (101,102). Melanin absorbs UVB radiation, and therefore reduces MSH and ACTH levels and activity. MSH promotes melanocyte proliferation and survival (103)

#### Stomach parietal cells

These cells secrete acid into the stomach under control of the hormone gastrin. Gastrin is secreted by stomach cells when pH is not acidic enough. Gastrin is the main growth factor for stomach parietal cells (104).

#### Liver cholangiocytes

Cholangiocytes form the bile ducts in the liver. They secrete water and bicarbonate under control of the hormone secretin. This hormone is secreted by cells in the duodenum when pH is acidic. Secretin production is inhibited by bicarbonate in the bile which reduces acidity in the duodenum. Secretin is the main growth factor for cholangiocytes (105,106)

#### Renin secreting cells

The juxtaglomerular cells in the kidney secrete renin that regulates blood pressure through a hormone cascade including angiotensin. Renin secretion by juxtaglomerular cells is inhibited by high blood pressure (107). High blood pressure was also suggested to inhibit the proliferation of these cells (108,109). For example, in “Goldblatt kidneys” patients, the adjacent blood vessels are partially blocked, causing the juxtaglomerular cells to sense a locally low blood flow. This results in increased renin secretion leading to high systemic blood pressure (110). Turgeon and Sommers found hyperplasia of juxtaglomerular cells in these patients, which was inversely correlated with hypertension duration.

#### Pituitary corticotophs

Corticotroph cells secrete ACTH and other POMC peptides, when activated by CRH secreted by the hypothalamus. CRH is the main growth factor for corticotrophs (111,112). ACTH inhibits CRH by an indirect path: ACTH activates the adrenal cortex to secrete cortisol, which shuts down CRH (and ACTH) production (98). There is also evidence for a direct inhibition of CRH secretion by ACTH in what is called a short feedback loop (113).

#### Pituitary somatotrophs

Somatotroph cells secrete growth hormone (GH) when activated by GHRH secreted by the hypothalamus (114). GHRH is the main growth factor for somatotrophs (115). GH inhibits GHRH secretion by an indirect path: GH activates the liver to secrete IGF1, which shuts down GHRH (and GH) production (114).

#### Pancreatic alpha cells

Alpha cells in the pancreas secrete the hormone glucagon, which activates glucose production in the liver. The regulation of secretion and proliferation in these cells is complex and a subject of current research. Glucagon secretion is inhibited by glucose, and glucagon is a growth factor for alpha cells (41). If glucagon acts in a local autocrine manner, this circuit is prone to mutant expansion of glucagon hyper-secreting cells, including glucose hypo-sensing alpha cells. Such cells secrete more glucagon and due to the autocrine effect, have higher growth rate. There are additional regulatory steps making the circuit more complex, and prohibiting a decisive analysis at this stage. For example, insulin also plays a role in glucagon secretion and alpha cell proliferation (116).

### S3: ASHM model with relative killing rate

We present a version of the ASHM model in which T cells killing is according to the level of antigen in a cell relative to the mean antigen level in the tissue. Such mechanisms have been theoretically proposed by Sontag (2017), based on earlier work (117,118).

In this case, ASHM removal goes as 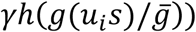 where 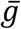 is the average antigen level 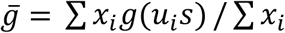. Thus,

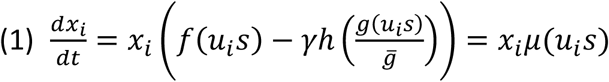

The condition to remove any mis-sensing mutant (any *u*_*i*_) on a background of wild-type cells (*u*_*i*_ = 1), is

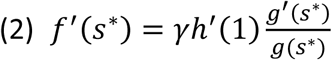

For linear secretion and growth functions, *g* = *g*_0_*u*_*i*_*s, f* = *μ*_0_(*u*_*i*_ *s* − *s*_0_), and Hill-like killing with cooperativity 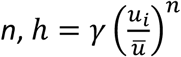, this condition is *γ*/*μ*_0_ *s*_*o*_ = 1/(*n* − 1).

To model the effect of mutant expansion on homeostasis, we again use the model of Karin et al (18). For linear secretion and growth functions and Hill-like ASHM killing, the wild-type set point is 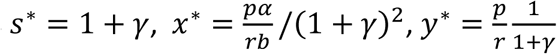. Thus the ASHM strength *γ* affects the set-point of the signal and cells. In autoimmune disease (large *γ*), tissue is lost and the signal loses homeostasis and.

In this case, the model has a new prediction: in autoimmune disease, the tissue will not be wholly destroyed, but will consist of hypo-secreting cells. These are variants or mutants that proliferate slowly and secrete poorly. They evade immune attack due to the differential sensing, and can persist as a small population. The larger immune killing rate gamma, the smaller this population.

Such a population of hypo-secreting cells are observed in late stages of T1D in mice (119,120). They have been described as “protected from the immune system” (121). This may be explained by the differential recognition in the ASHM model.

